# Scalable Classification of Organisms into a Taxonomy Using Hierarchical Supervised Learners

**DOI:** 10.1101/2020.02.04.933374

**Authors:** Gihad N. Sohsah, Ali Reza Ibrahimzada, Huzeyfe Ayaz, Ali Cakmak

## Abstract

Taxonomy of living organisms gains major importance in making the study of vastly heterogeneous living things easier. In addition, various fields of applied biology (e.g., agriculture) depend on classification of living creatures. Specific fragments of the DNA sequence of a living organism have been defined as DNA barcodes and can be used as markers to identify species efficiently and effectively. The existing DNA barcode-based classification approaches suffer from three major issues: (i) most of them assume that the classification is done within a given taxonomic class and/or input sequences are prealigned, (ii) highly performing classifiers, such as SVM, cannot scale to large taxonomies due to high memory requirements, (iii) mutations and noise in input DNA sequences greatly reduce the taxonomic classification accuracy. In order to address these issues, we propose a multi-level hierarchical classifier framework to automatically assign taxonomy labels to DNA sequences. We utilize an alignment-free approach called spectrum kernel method for feature extraction. We build a proof-of-concept hierarchical classifier with two levels, and evaluated it on real DNA sequence data from BOLD systems. We demonstrate that the proposed framework provides higher accuracy than regular classifiers. Besides, hierarchical framework scales better to large datasets enabling researchers to employ classifiers with high accuracy and high memory requirement on large datasets. Furthermore, we show that the proposed framework is more robust to mutations and noise in sequence data than the non-hierarchical classifiers.

## 1. Introduction

Classification of living organisms is a key problem in both biology and computer science. This is a result of being at the core of many fields of applied biology such as agriculture and public health. Using traditional morphological keys for classification is often efficient only for a particular gender or life stage. Besides, this method is slow and expensive, as it requires the time and effort of highly experienced taxonomists.

DNA Barcoding has gained significant attention [1, 2] in the scientific community after it was firstly introduced by Herbert et al. [3]. Specific gene regions have been chosen as markers that can distinguish between different species [4–6]. For animal groups, cytochrome c oxidase 1 gene (COI) is used as a barcode, while matK and rbcL are used for identifying land plants, and ITS is used for fungi [7]. The problem then translates into classifying barcodes to a known species in a fast and efficient way [8].

There is a number of methods [9, 17] that tackle with the DNA barcode-based classification problem using the tools of sequence comparison and alignment. However, aligning multiple sequences in an optimal way is computationally costly. Using alignment-free kernel methods has been proven efficient for this problem [18]. In this approach, the occurrences of each possible fixed-length substring are counted in each DNA barcode sequence. These substrings are called *k-mers*, where *k* is an integer parameter that corresponds to the length of the substring. These *k-mers* are then used as features with their corresponding count per sequence as feature values.

Weitschek et al. [9] propose and compare several machine learning classification techniques to determine species given a DNA Barcode. In their work, they include several machine learning techniques like Support Vector Machines [10], the rule-based method RIPPER [11], the decision tree C4.5 [12], and the Naïve Bayes [13]. A major drawback is that they consider only pre-aligned sequences. Besides, they perform the classification strictly within the scope of specific taxonomic classes like bats, birds, fungi, or fishes. Hence, in order to classify a given barcode sequence, it first needs to be aligned, and one also needs to know to which taxonomic class the given sequence belongs to.

Austerlitz et al. [14] compare phylogenetic and statistical classification methods using datasets that included six different genera. They consider Neighbor Joining [15] and PHYML [16] as phylogenetic methods, and k-nearest neighbor, classification and regression trees (CART), random forests, and support vector machine as statistical classification methods. However, in their work, they assume apriori knowledge about the genus of the sequence and they use pre-aligned sequences. In this work, the method that obtained the best score varied according to the dataset. However, the support vector machine method attained the best score at three out of six datasets while each of the other methods achieved the best score for at most two datasets.

Another work by Weitschek et al. [17] exploits a supervised machine learning approach that selects suitable nucleotide positions and computes the logic formulas for species classification. However, similar to the above work, it also requires prealigned DNA barcode sequences.

Kuksa and Pavlovic [18] employ an alignment-free method based on *k-mers* for DNA barcode classification and analytics. They exploit *10-mer* features to train classification models using two classes of algorithms: Nearest Neighbor and SVM.

Zhao et al. [19] proposes the use of a new set of classification features that are based on covariance of nucleotides in DNA barcodes. The computed features are later exploited in a random forest classifier to perform phylogenetic analysis on a particular fungi species.

Kabir et al. [20] explores the power of different supervised learning methods for DNA barcode-based classification. To this end, they evaluate different classifiers including Simple Logistic Function [21], (ii) IBk from Lazy classifier [22], (iii) PART from Rule based classifier [23], (iv) Random Forest from Tree based classifier [24], (v) Attribute Selected Classifier, and (vi) Bagging from Meta classifiers [25].

In most of the existing studies that focus on the problem of organism classification using DNA sequences, the classification is mainly performed within a specific taxonomic class assuming apriori knowledge about the given to-be-classified sequence. This assumption may not always hold true, e.g., when inspecting fossil remains or sequences extracted from mud-samples and earth layers. In such situations, it is hard to identify whether these sequences belong to the class of bats, birds, rodents, fishes, etc. Furthermore, highly performing classifiers, such as SVM, cannot scale to large taxonomies due to high memory requirements. Besides, mutations and noise in input DNA sequences greatly reduce the taxonomic classification.

In order to address the above issues, in this paper, we introduce a hierarchical framework that can be extended into one hierarchical classifier capable of classifying any DNA barcode sequence without any apriori knowledge about its taxonomic tree. This framework utilizes Support Vector Classifiers in order to build a two-stage classifier that can predict the species given the DNA barcode sequence only, without the need to compute any sequence alignment. Our framework enables leveraging the strength of the Support Vector Classifiers while overcoming the scalability issues that arise when the number of classes increases or when the data matrix size grows.

In order to establish a proof-of-concept, we test the proposed approach on three different datasets obtained from BOLD Systems [26] under the Chordata taxonomic class: (1) Aves, which is the dataset of the birds. (2) Chiroptera, the bats dataset. 3) Rodentia, the dataset of the rodents. For each dataset, classification accuracies of different classifiers using different subsequence lengths *k* during feature extraction are compared. We observe that the SVM classifier with a linear kernel outperforms all the other methods for larger *k* values. The Random Forest classifier, on the other hand, outperforms the SVM-based classifiers when *k* is relatively small. Then, we merge the three datasets to examine the scalability of each classification method. For larger datasets, SVM was not a feasible solution, since the data matrix was not representable. Nevertheless, the RF method worked with a test accuracy of 91.1%. In order to overcome this scalability drawback and utilize the high accuracy of the linear kernel SVM, we build a hierarchical SVM-based classifier, and demonstrate that it outperforms the non-hierarchical regular classifier (95.9% vs. 91.1%). Besides, we also study the robustness of the proposed method by introducing artificial mutations to the sequences with increasing ratios. Our experimental results show that as mutations rates increase, the proposed hierarchical classifier framework exhibits more robustness than the other non-hierarchical classifiers.

Our contributions in this paper are as follows:

- We propose a multi-level hierarchical DNA sequence classification framework, and build a proof-of-concept instance with two taxonomic levels. The proposed framework can be extended to predict the species within larger scopes that go beyond just the taxonomic class level. More specifically, it can be used as a blueprint in building a full supervised classifier that can classify all life forms.
- We demonstrate that the hierarchical classifier framework classifies DNA sequences with higher accuracy than the regular stand-alone classifiers.
- The proposed framework allows taking advantage of SVM’s high accuracy prediction power for larger datasets as well by increasing its scalability with multi-level organization. While regular SVM-based classifiers run out of memory when trained on a large dataset on a decently configured test environment, the hierarchical SVM-based classifier successfully runs on the same test hardware and dataset.
- We demonstrate that with hierarchical classifier, the robustness of classification in the presence of mutations and/or noise in sequence data is higher than the regular classifiers.

## 2. Methods

### 2.1. Kernel-based alignment-free method for feature extraction

Kernel-based methods are employed to represent sequences with variable lengths and also to avoid the burden of handling insertions and deletions. Kernel-based methods have been proven to be efficient in a number of similar tasks like protein-protein interaction prediction and protein classification [27–29]. They have been also demonstrated to be effective in tackling the problem of species classification using DNA barcodes [18]. In this method, sequences are represented as collections of short substring kernels of length *k*. These substrings are called *k-mers*. Figure 1 illustrates how a sequence can be represented as a vector of *k-mers* frequencies, where *k* = 5.

**Fig. 1:**
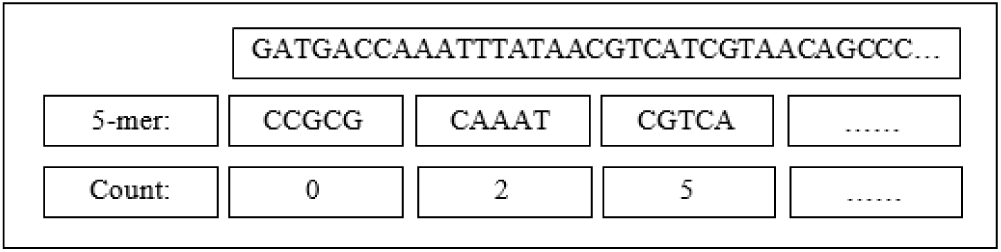
An example of how 5-mer kernels are used to represent a DNA barcode sequence.

The number of k-mers increases exponentially with *k*. Since we have 4 bases (i.e., A, C, G, T), the number of all *k-mers* is 4^*k*^. The occurrence frequency of these *k-mers* are then used as features. A variation of this method employs mismatch-kernels for feature extraction [29,31]. In this case, at most *m* mismatches are allowed within a substring. This can enhance the results of the classification task by making the data matrix denser which is desirable for most of the classification algorithms. However, in this paper, we do not study the effect of allowing mismatches and the effect of changing the value of *m*. Instead, we focus on building a hierarchical classifier that allows the classification tasks to be more efficiently performed on large datasets that include different taxonomic classes.

### 2.2. Scalable Supervised Learning

In most of the related work, it has been shown that it is possible to train a supervised classifier that has the ability to predict the species given the DNA barcode sequence. However, there were two factors that kept the effectiveness of such approaches restricted. First, these studies carried out the prediction effort within a specific taxonomic organism rank, e.g., performing the experiments on the taxonomic class level such as Chiroptera, Rodentia, Aves, Mammalia, etc. [9, 18], or on the taxonomic genus level as in [14]. Hence, they assume the availability of apriori knowledge of the taxonomic class or genus to which a specimen or a sequence belong to. Such an assumption may not hold true in many cases, such as when inspecting samples from a lake or soil.

Second, among supervised classification algorithms, Support Vector Machines (SVM) are commonly employed in taxonomy classification, as it provides more accurate results than many other methods. However, SVM suffers from scalability issues when the number of classes in the dataset increases, as the data matrix grows in size. All these reasons hinder its use as an efficient classification algorithm to train a classifier that predicts the species directly from the DNA barcode sequence. As, in that case, the number of classes would be the number of all known species, and the dataset would be all the data samples available on BOLD Systems. Motivated by the above observations, in this paper, we propose a two-stage hierarchical classifier inspired by the hierarchical nature of the taxonomy tree. The first stage predicts the taxonomic class. Then, according to the prediction of the first-stage classifier, the feature vector representing a given DNA barcode sequence is directed to the corresponding classifier trained to predict the species within that taxonomic class. A diagram of this framework is shown in Figure 2.

**Fig. 2:**
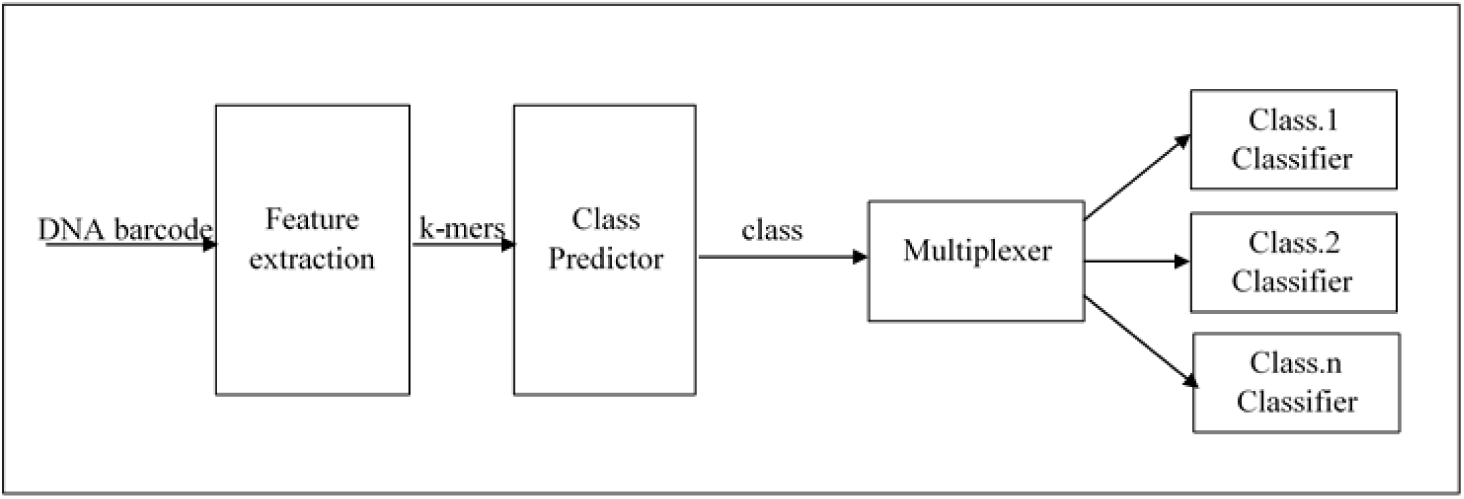
Two-stage hierarchical classifier for predicting the species without any apriori knowledge about its taxonomic class

The illustrated framework is used to train a classifier capable of predicting the species name for a given DNA barcode sequence out of 1090 species appeared in the datasets used in this work. Although results were obtained for 1090 species that belong to three different taxonomic classes (Aves, Rodentia, and Chiroptera), the extension of this work into one hierarchical learner capable of classifying all known living things is straight forward.

## 3. Results

In this section, the proposed framework is experimentally assessed in terms of accuracy, scalability, and robustness. We also compare it to regular non-hierarchical approaches. As a proof-of concept, two popular supervised learning classifiers are contrasted, namely, SVM and Random Forests. We study how different classification methods perform and how varying *k* affects their performance. In the first subsection, the performance within the scope of a taxonomic class level is evaluated using three different datasets (Rodentia, Aves, and Chiroptera). Then, the robustness aspects of the classifiers are studied in the presence of mutations or noise in data. Finally, a scalability analysis of the proposed hierarchical classification is performed in comparison with the non-hierarchical classification.

All the experiments for this paper were carried out on a DELL R720 server whose specifications are summarized in Table 1. All the scripts are coded in *Python* using *Scikit - Learn* machine learning library to implement SVM [32] and Random Forest [33] classifiers. Besides, *Matplotlib* and *Seaborn* are used for the visualization.

**Table 1:** Hardware Specification

| Instance Type | vCPU | Mem (GiB) | Storage (TB) |
| --- | --- | --- | --- |
| DELL R720 | 24 | 80 | 2.4 |

### 3.1. Datasets

For this study, the datasets are obtained from Barcode of Life Data (BOLD Systems) [26] which is an initiative to support the generation and application of DNA barcode data. It contains 4,212,621 DNA barcode sequences for animals, plants, fungi, and all other life forms. The specimen is collected from different sites by different organizations worldwide. Through the portal of BOLD Systems, data for different life forms may be downloaded in various formats including XML and tab-separated text.

In this paper, three datasets were used, Chiroptera dataset, Aves dataset, and Rodentia dataset. As a preprocessing step, all the sequences that are less than 657 in length were removed since the full length of the COI segment used as a DNA barcode sequence is 657 bp [30]. However, sequences with ambiguous letters like “Ns” and dashes “-” were kept. Table 2 gives a summary of datasets after the preprocessing step.

**Table 2:** Class Datasets Summary

| Dataset | No. of species | No. of samples |
| --- | --- | --- |
| Chiroptera | 122 | 4731 |
| Rodentia | 127 | 3653 |
| Aves | 841 | 4192 |

To gain insights about the distribution of the datasets, the distribution of each dataset is plotted in Figures 3, 4, 5. Furthermore, the maximum, minimum, and average frequencies were calculated and reported in Table 3.

**Table 3:** Class Datasets Species Frequencies Summary

| Dataset | Maximum frequency | Minimum frequency | Average frequency |
| --- | --- | --- | --- |
| Chiroptera | 0.2 | 0.0002 | 0.008 |
| Rodentia | 0.1 | 0.0002 | 0.008 |
| Aves | 0.02 | 0.0002 | 0.001 |

**Fig. 3:**
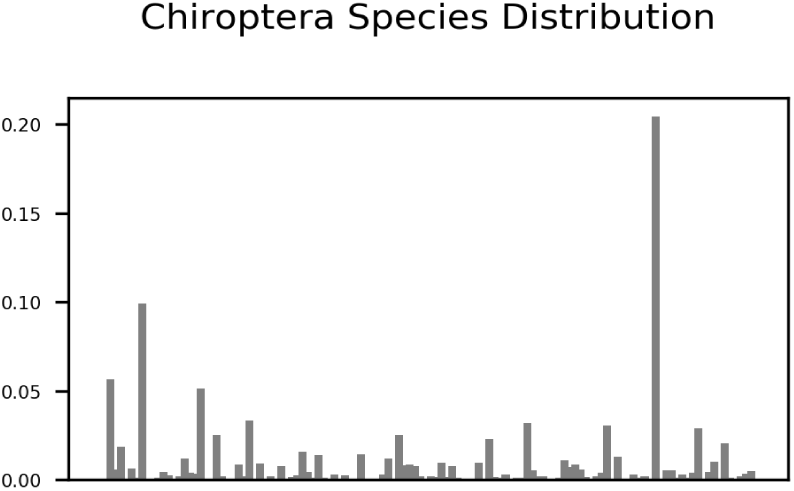
The distribution of all species in Chiroptera dataset.

**Fig. 4:**
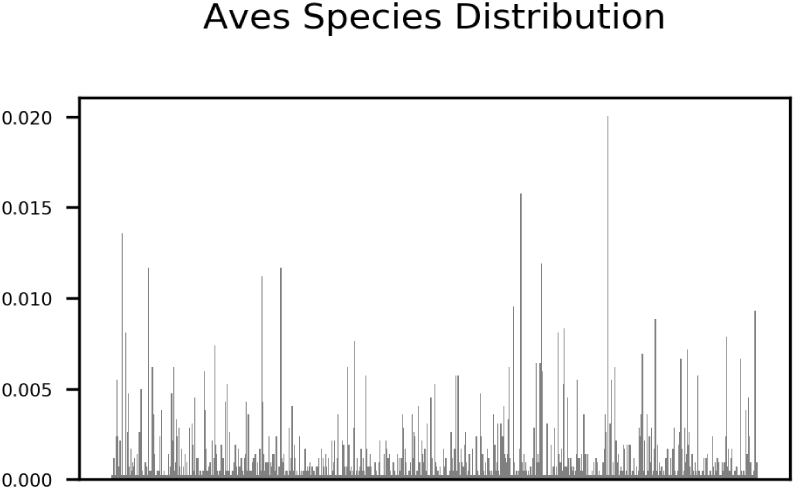
The distribution of all species in Aves dataset.

**Fig. 5:**
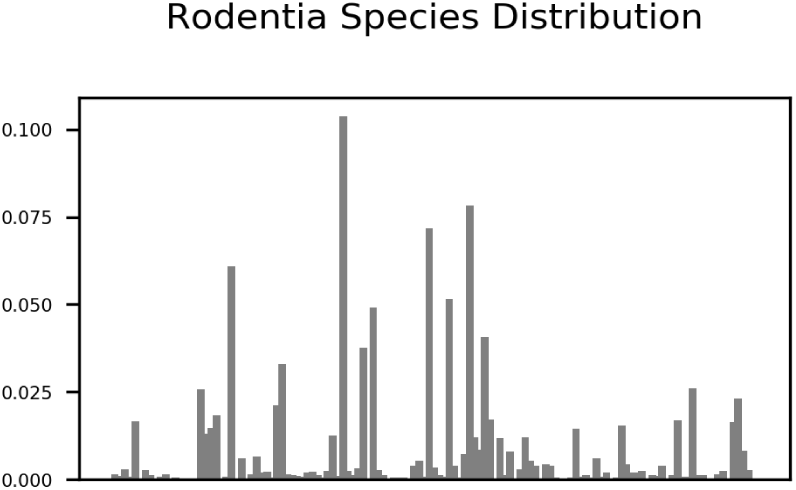
The distribution of all species in Rodentia dataset.

Before each experiment, datasets are randomized and partitioned into two splits, a training set (90% of the samples) using which a model is trained, and a test set (10% of the samples) on which the models were evaluated. The models are evaluated using 10-folds cross validation by repeating the dataset randomization, splitting, training, and testing steps 10 times, and the resulting accuracy values are averaged. Moreover, in order to make sure that class and species distributions match in both training and test splits, stratified sampling is applied during 10-fold cross validation. Here, we report the averages over all runs.

### 3.2. Accuracy Evaluation of Non-hierarchical Classifiers

In this part of our experiments, the effect of changing the length of the subsequence kernel *k-mers* on the accuracy of non-hierarchical classifiers is studied. To this end, using each of the datasets, three different classifiers (i.e., Random Forest with 10 estimators, SVM with linear kernel, and SVM with radial kernel) are trained and tested using 10-folds cross-validation. In all experiments, for SVM with radial kernel, the kernel width is set to the number of samples in the training split at each iteration. In all charts, boxplots are also included in order to demonstrate the variability of accuracy values across different folds during cross validation. Please note that in order to prevent the overlap among box plots as well as the original accuracy lines, box plots are slightly shifted so that they do not block each other in the visualization.

#### 3.2.1. Chiroptera Dataset

The results for the Chiroptera dataset are visualized in Figure 6. The SVM classifier with a linear kernel, for larger *k* values, outperforms the other classification methods, while the Random Forest classifier performs better for lower *k* values. It can also be noted that the accuracy values obtained with the SVM classifier with a linear kernel increase as *k* increases in the range [1, 7] without any drop, unlike the SVM classifier with a radial kernel which experiences accuracy drop for larger values of k.

**Fig. 6:**
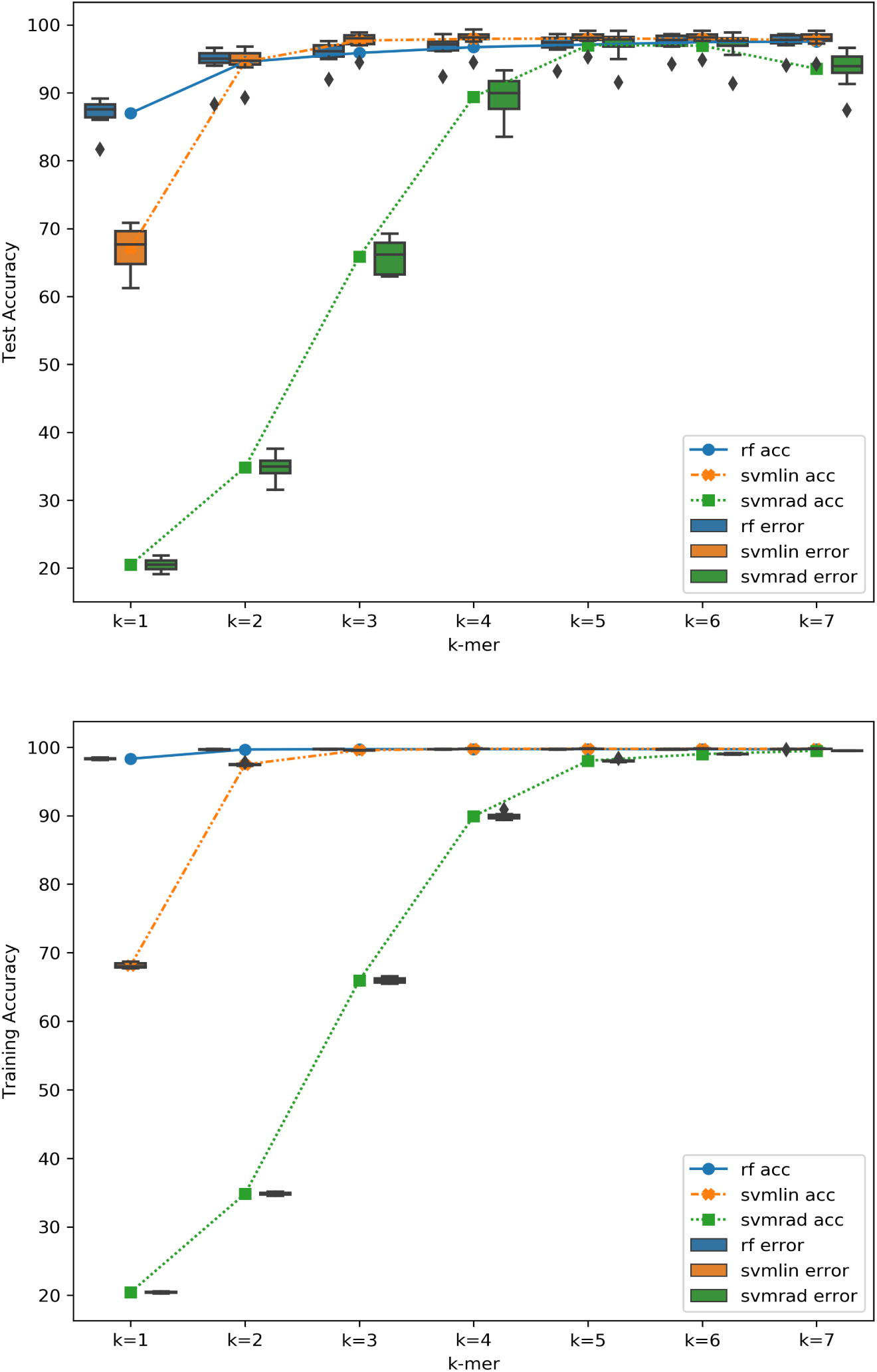
The effect of changing the value of *k* on the classification accuracies of three different classification methods (Random Forest with 10 estimators, SVM with linear kernel, and SVM with radial kernel) trained and tested on the Chiroptera dataset.

#### 3.2.2. Rodentia Dataset

Similar observations are made for the Rodentia dataset as illustrated in Figure 7.

**Fig. 7:**
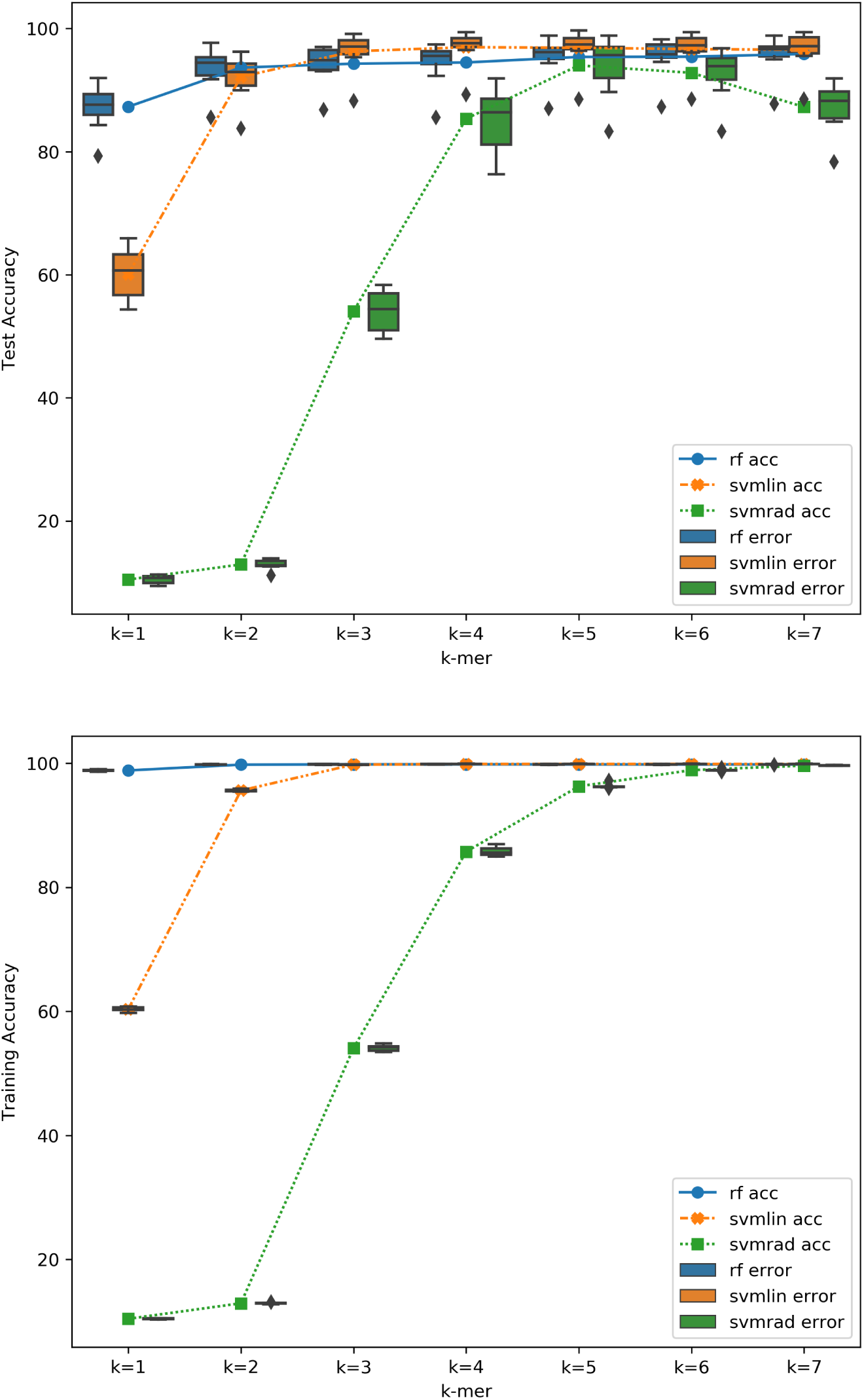
The effect of changing the value of *k* on the classification accuracies of three different classification methods (Random Forest with 10 estimators, SVM-linear kernel, and SVM-radial kernel) trained and tested on the Rodentia dataset.

#### 3.2.3. Aves Dataset

As for the Aves dataset, although the test accuracy is lower than the above two datasets (see Table 4), the relative rank and behavior of the classifiers are similar to the above results with the change of *k*, as shown in Figure 8. It can be concluded that, as long as the memory resources allow larger values of *k*, it is possible to train an SVM classifier with a linear kernel that achieves better classification accuracies than both an SVM classifier with a radial kernel and an RF classifier. On the other hand, if the memory limitations hindered the increase of *k*, one may opt for Random Forest classifier, as we show in the next section that random forest requires less memory.

**Table 4:** Maximum Accuracies Obtained for Each Dataset

| Dataset | training accuracy | test accuracy |
| --- | --- | --- |
| Chiroptera | 99.7% | 98% |
| Rodentia | 99.9% | 97% |
| Aves | 99.9% | 88% |

**Fig. 8:**
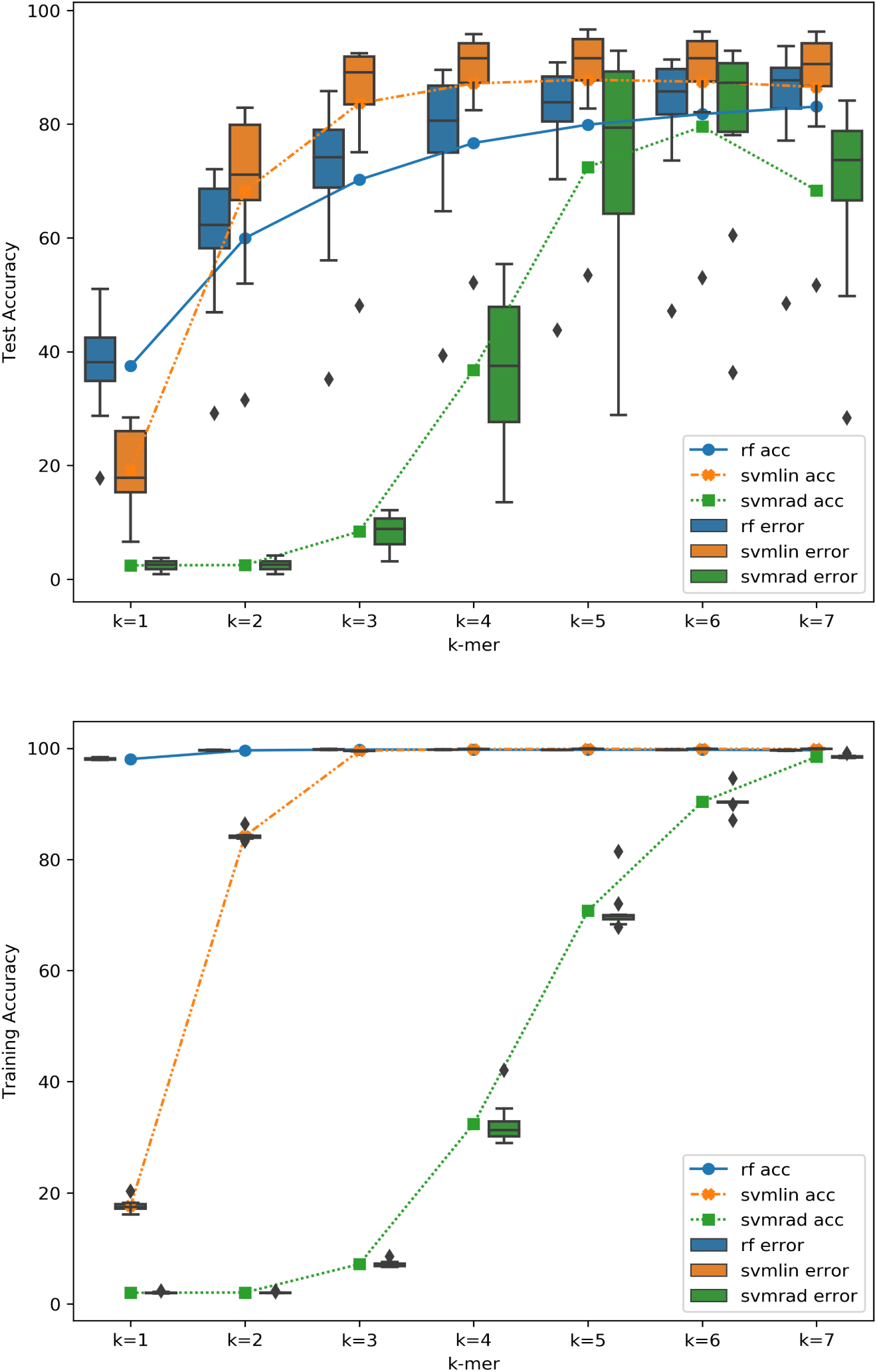
The effect of changing the value of *k* on the classification accuracies of three different classification methods (Random Forest with 10 estimators, SVM with linear kernel, and SVM with radial kernel) trained and tested on the Aves dataset.

To sum up, the best accuracies for all datasets are provided by an SVM classifier with a linear kernel trained using a subsequence of length *k* = 7. As Table 4 shows, the classification accuracy for the Aves dataset is comparably low. The reason behind this is mainly the insufficient number of samples per species. Since, as summarized in Table 2, the number of species in the Aves dataset is 841 which is roughly 7 times more than the number of species in the other two datasets, while the number of samples are almost the same in each dataset.

### 3.3. Scalability of Non-hierarchical Classifiers

In order to study how efficiently different classification methods scale to larger datasets, the three datasets used in this paper, Chiroptera, Rodentia, and Aves datasets, are merged into a single dataset. Then, the above three classifiers are trained using the same settings used in the previous section. Unfortunately, the attempts to train the SVM classifiers (with both linear and radial kernels) failed due to memory limitations, despite the decent memory size of the test machine. However, the Random Forest classifier was trained successfully and provided the results illustrated in Figure 9. The maximum test accuracy (91.1% approx.) was obtained at *k* = 7.

**Fig. 9:**
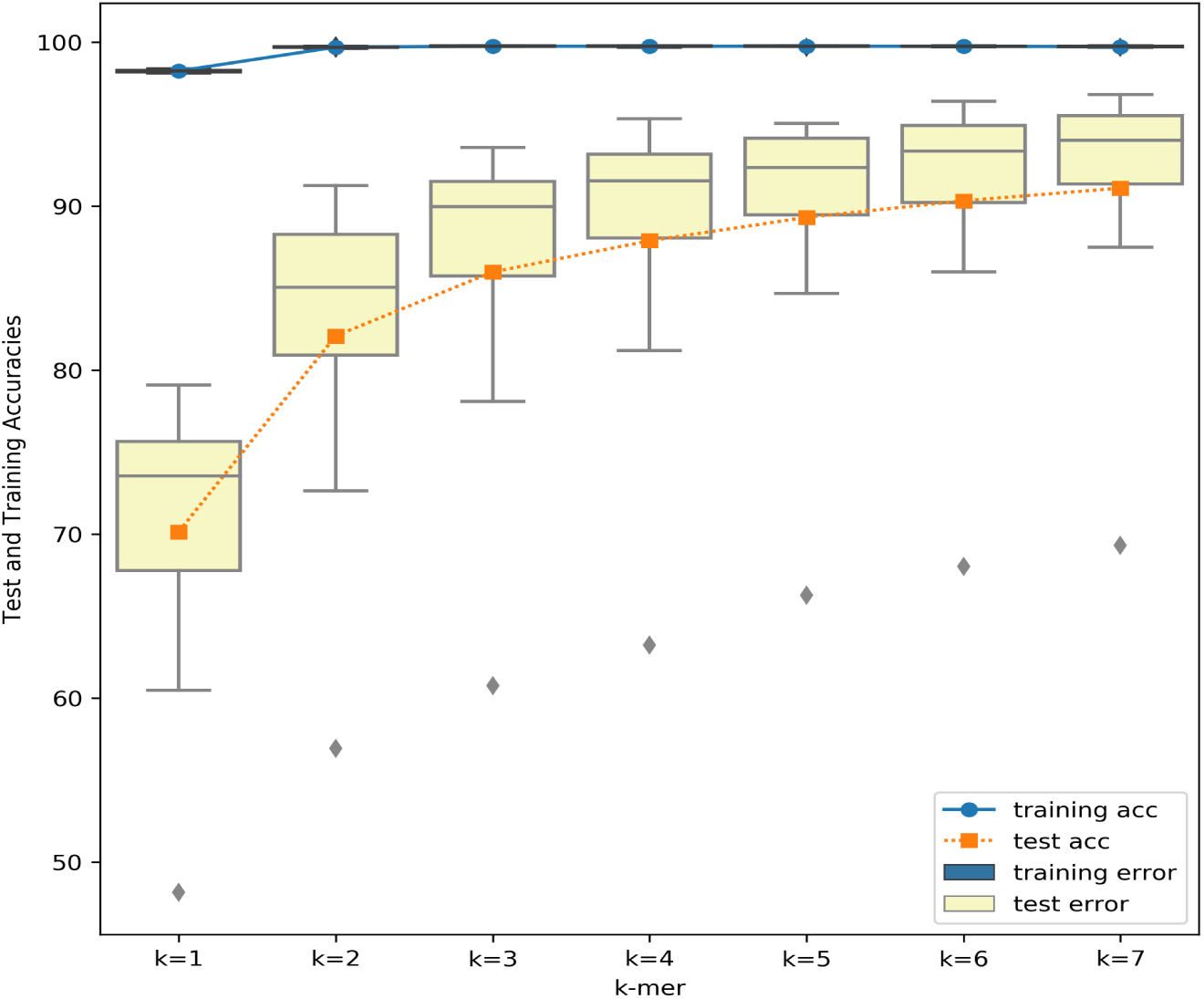
The effect of changing the value of *k* on the classification accuracies of a Random Forest Classifier with 10 estimators trained and tested on all the three datasets merged together.

In order to be able to leverage the strength of SVM classifiers, while overcoming their scalability issues, we employ our proposed hierarchical framework as demonstrated next.

### 3.4. Accuracy and Scalability of the Hierarchical Classification Framework

#### 3.4.1. Taxonomic Class Predictor

The first stage of the proposed framework (see Figure 2) involves the training of a taxonomic class predictor. On the merged dataset (that involves Chiroptera, Aves, and Rodentia datasets), three different taxonomic class classifiers are trained to predict whether a DNA barcode sequence belongs to the Rodentia class, Chiroptera class, or Aves class. The accuracies of these three classifiers with varying subsequence length *k* are presented in Figure 10.

**Fig. 10:**
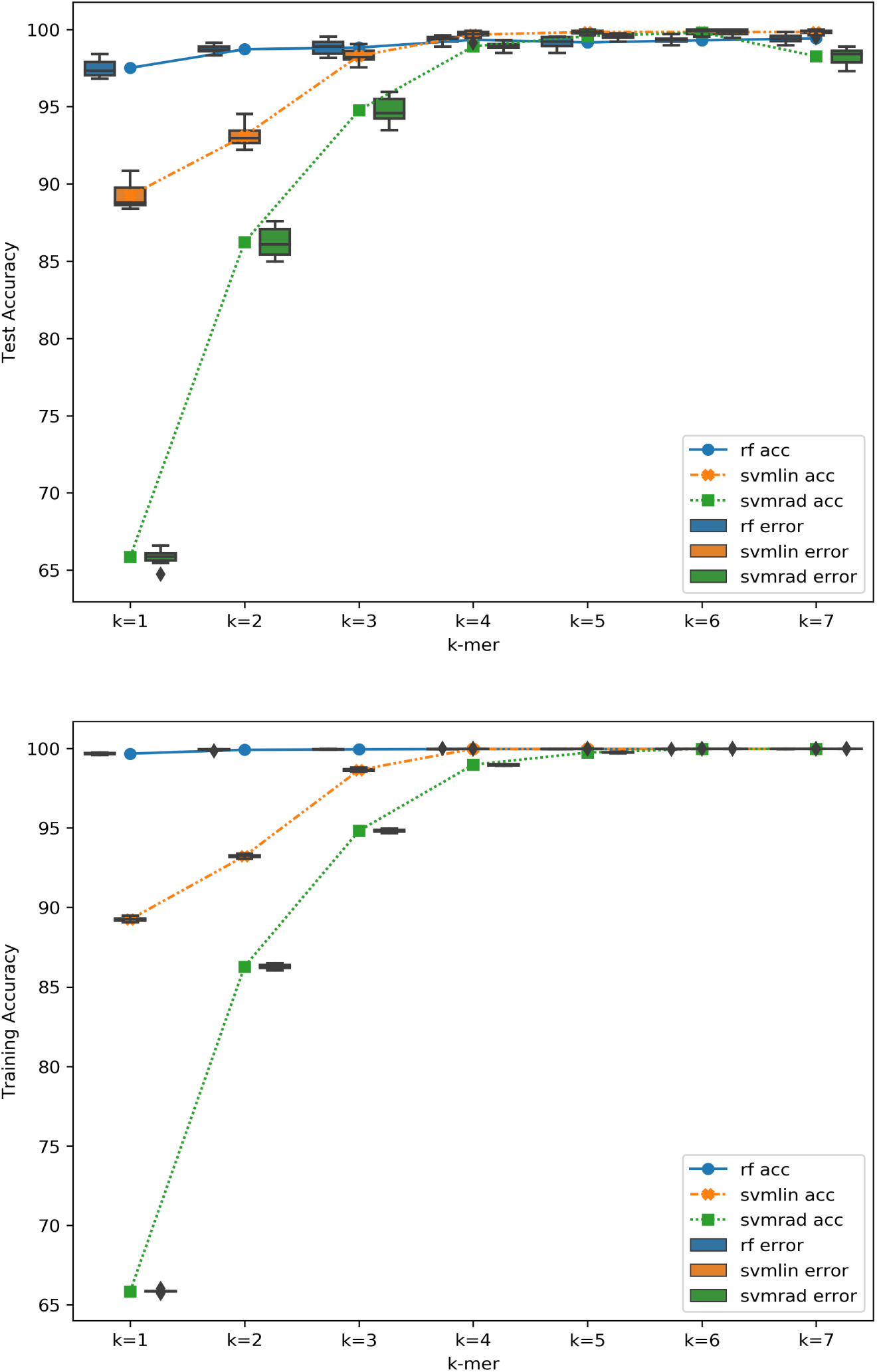
The effect of changing the value of *k* on the accuracies of three different classification methods (Random Forest with 10 estimators, SVM with linear kernel, and SVM with radial kernel) trained to predict the taxonomic class on the merged dataset.

From Figure 10, we observe that the best accuracies are obtained when using SVM classifier with a linear kernel and setting the subsequence length *k* to be 7. The test accuracy in this case is 99.9% which means that we are able to predict the taxonomic class with an error of 0.1%, and then pass the sequence to a class-based classifier capable of predicting the species with the accuracies given by Table 4.

In the hierarchical classifier, two SVM classifiers with linear kernels are combined in a hierarchical manner. The first one assigns a taxonomic class to each DNA barcode sequence, and passes the sequence down to the second level species classifier to assign a species. The framework for the hierarchical method is illustrated in Methods section. SVM classifiers with linear kernels are chosen due to their relatively high performance as shown with the above experimental results. Figure 11 shows how the performance of the two-stages SVM-hierarchical classifier changes with the value of *k*. Table 5 also compares the accuracy of the non-hierarchical classifier (with the RF classifier) trained on the merged dataset (with k = 7) against the accuracy of the hierarchical classifier on the same dataset.

The above results demonstrate that the proposed hierarchical classification framework provide superior accuracy performance than the non-hierarchical classifier. It also overcomes the memory limitation that is discussed above.

#### 3.4.2. Putting It Altogether: Hierarchical Classifier

**Fig. 11:**
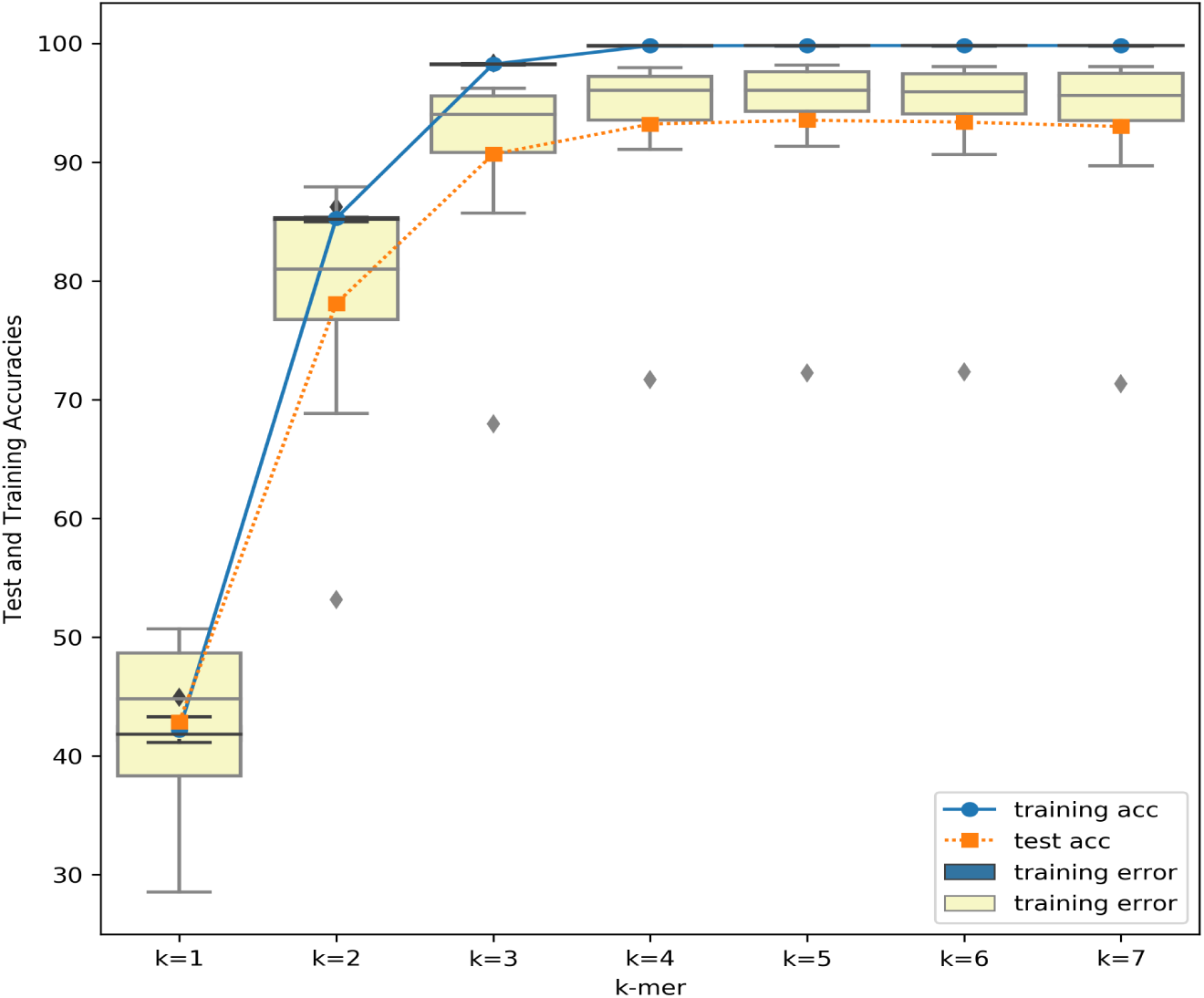
The effect of changing the value of *k* on the accuracy of the hierarchical classifier that employs linear kernel SVM sub-classifiers.

**Table 5:** Comparison of Methods

| Method | training accuracy | test accuracy |
| --- | --- | --- |
| Random Forest (10 estimators) | 99.5% | 88.5% |
| Hierarchical SVM | 99.8% | 95.9% |

### 3.5. Robustness Analysis

In this section, we study the robustness of hierarchical and non-hierarchical classification frameworks in the presence of mutations and/or noise in DNA sequences. In order to simulate the mutations or sequencing noise, we randomly introduce artificial mutations in the DNA barcode sequences with different ratios. More specifically, the mutation ratios are varied in the range [0,1] with a step of 0.1, and the accuracy for each classifier is reported. For each mutation ratio, the number of mutation positions is calculated by multiplying the ratio by the sequence length, and then that many mutations are introduced at random positions. Replacement characters are chosen randomly from the set -, A, G, C, T.

As discussed in the above experiments, one single SVM classifier could not be trained using the merged dataset due to high memory requirements. However, the random forest algorithm could scale to train one classifier capable of predicting the species for a given DNA barcode sequence regardless of the taxonomic class in the merged dataset. Here, we compare the robustness of the proposed hierarchical classification framework (with SVM and RF subclassifiers, separately) to that of non-hierarchical classifier (built with RF). The kernel that is employed in all classifiers is a linear kernel due to the relative efficiency of the SVM-linear classifiers as illustrated by figures 6, 7, and 8. All studied classifiers are trained and tested with 10-folds cross validation using the combined dataset that merges the three taxonomic classes considered in this work.

Figures 12 and 13 present the test and the training accuracies of classifiers, respectively. These results demonstrate that the proposed hierarchical framework provides more robust accuracy performance in the existence of mutations of noise in data in comparison to the conventional non-hierarchical structure.

**Fig. 12:**
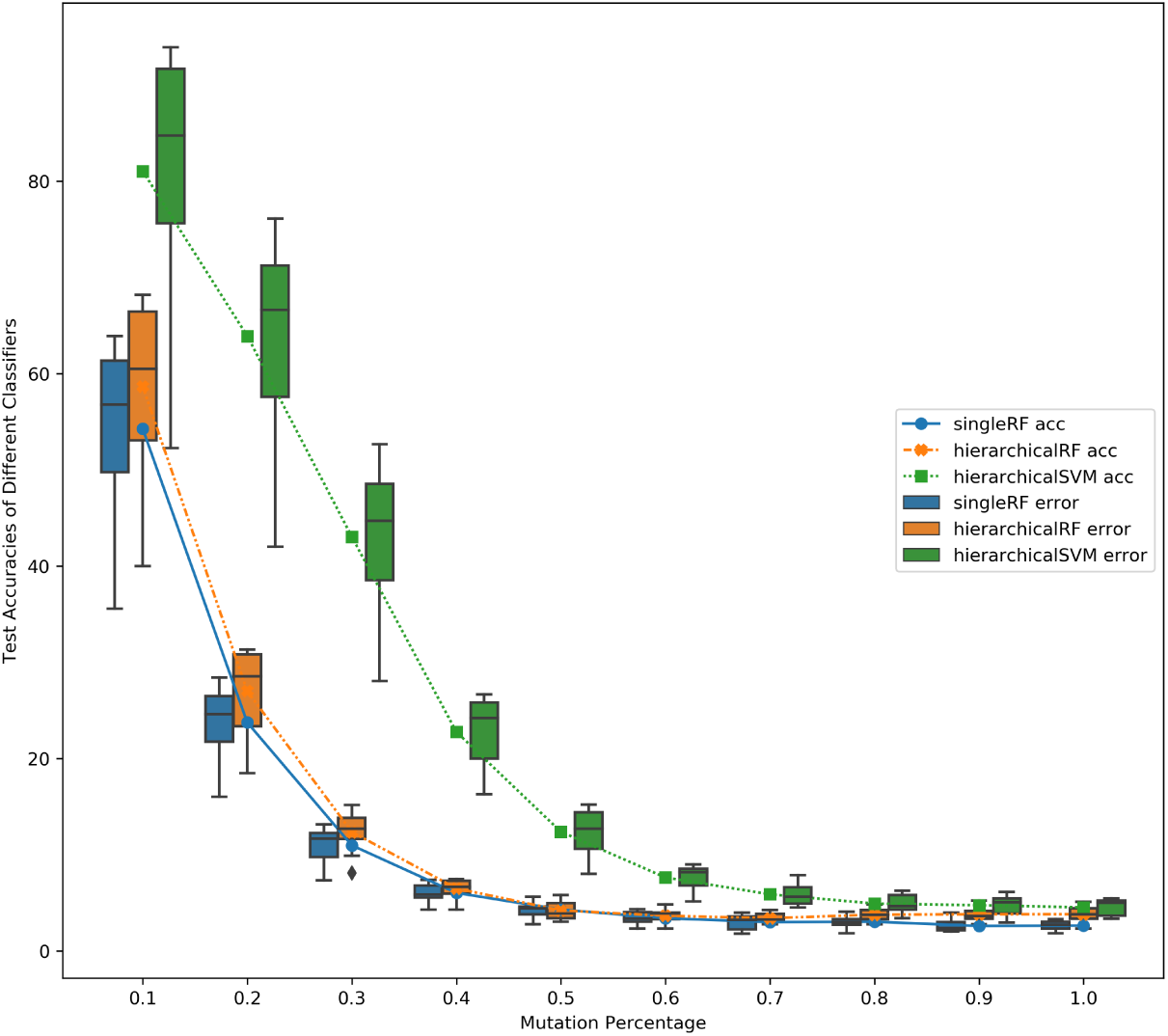
The effect of introducing artificial mutations on the classification test accuracies of a hierarchical RF classifier with the number of trees set to 10, non-hierarchical RF classifier with the number of trees set to 10, a hierarchical SVM classifier with linear kernel.

**Fig. 13:**
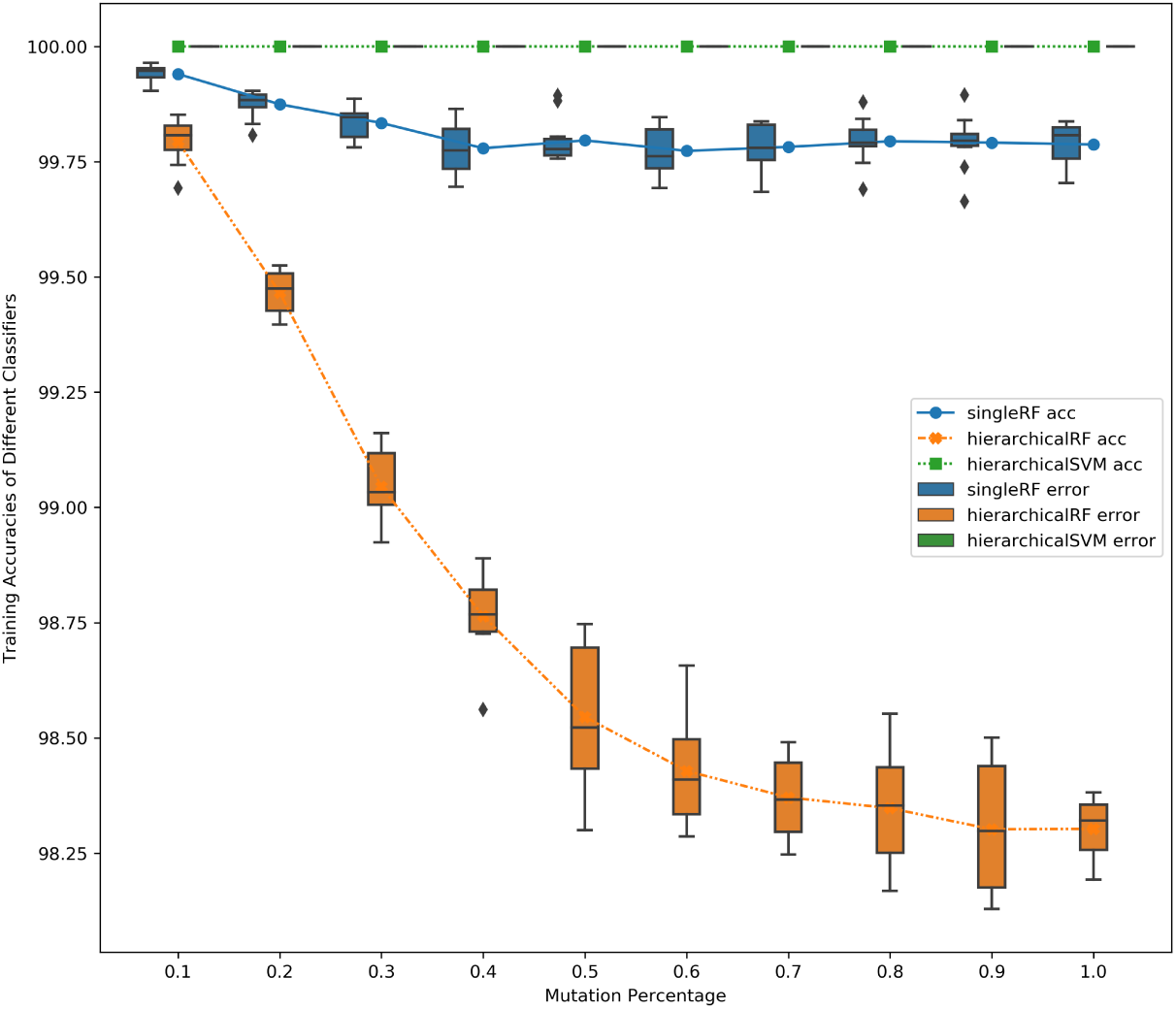
The corresponding training accuracies for the classifiers in the previous chart.

## 4. Conclusions

In this paper, the problem of assigning taxonomic labels to DNA sequences using supervised learning is studied. A multi-level hierarchical classification framework which combines multiple classifiers built for predicting a label (e.g., class, genus, species, etc.) at different levels in organism taxonomy is proposed. The proposed framework is evaluated on real data of 1090 species from BOLD systems. We demonstrate that, in comparison to the conventional supervised classifiers, the proposed method provides the following advantages: (i) better accuracy, (ii) improved scalability, (iii) more robustness against mutations or noise in sequence data.

## Authors’ information

***Gihad Sohsah*** received her Bachelor of Computer Engineering from Tanta University (Tanta, Egypt) in 2011. She has recently received her M.Sc. degree in Data Science from Istanbul Sehir University (Istanbul, Turkey). Gihad is mainly interested in both theoretical and applied Machine Learning and Artificial Intelligence.

***Ali Reza Ibrahimzada*** is currently studying his Bachelor in Computer Science and Engineering at Istanbul Sehir University (Istanbul, Turkey). Ali Reza is mainly interested in Machine Learning, Artificial Intelligence and Data Science. He is a member of Bioinformatics and Databases Lab at Istanbul Sehir University.

***Huzeyfe Ayaz*** is a sophomore student in Computer Science and Engineering department at Istanbul Sehir University (Istanbul, Turkey). He is interested in Data Science, Machine Learning, and AI. Currently, he works on several research projects to improve the public health within the Bioinformatics and Databases Lab at Istanbul Sehir University.

***Ali Cakmak*** received his B.Sc. degree in 2003 from the Computer Engineering Department at Bilkent University (Ankara, Turkey), and his Ph.D. degree in 2008 from the Electrical Engineering and Computer Science Department at Case Western Reserve University (Cleveland, OH). After completing his Ph.D., he worked as a post-doctoral research associate for a year in the same department. Then, he moved to the Silicon Valley, and worked as a senior software engineer as part of the Query Optimization Group at Oracle, Inc (Redwood Shores, CA). In Fall 2013, he joined Istanbul Sehir University (Istanbul, Turkey) as a faculty member in the Department of Computer Science. His research interests include bioinformatics, machine learning, data mining, databases, and data science. Dr. Cakmak is a recipient of TUBITAK (The Scientific and Technological Research Council of Turkey) Career Grant.

